# Molecular underpinning of *Hydra viridissima*-algal facultative symbiosis and vertical algal transmission

**DOI:** 10.64898/2026.05.21.726902

**Authors:** Joseph R. Tran, John Sittmann, Binbin Ma, Minshuai Zhu, Yixian Zheng, Minjie Hu

## Abstract

The symbiosis between algae and animals represents a relatively recent evolutionary innovation, and exemplified by species in the Cnidaria phylum. Cnidaria can harbor algae within a modified cellular organelle called the symbiosome in a process called endosymbiosis. This animal-algal symbiosis can be facultative or obligate. Algae acquisition occurs either by horizontal transmission, where free-swimming planula gain algae through feeding, or by algae deposition into the developing oocyte in vertical transmission[1–5]. Most studies focus on anthozoans that perform obligate endosymbiosis and transmit algae horizontally. How facultative endosymbiosis in combination with vertical algal transmission impacts the evolutionary adaptation between host and symbiont remains unclear. By studying *Hydra viridissima*, which performs facultative endosymbiosis and transmits its *Chlorella* algae vertically, we define different cell types and identify the endoderm cell lineage that gives rise to three major cell types hosting algae. Compared to obligate endosymbiosis[6, 7], *Hydra viridissima* algal host cells exhibit distinct features, including algal uptake, elevated oxidative phosphorylation and redox activities, and express genes that can provide ammonium for their algal symbionts. We further show when and how the developing *Hydra* oocytes may take up algae and where oocytes may obtain lipids. Since *Hydra* is amenable to genetic manipulations, our findings should enable mechanistic studies of how facultative endosymbiosis and vertical transmission evolve and adapt in a changing climate.

## Introduction

Some animal species have evolved a strategy to benefit from photosynthetic products by endosymbiotically-hosting algae. Most studies focus on the coral anthozoans that perform obligate endosymbiosis with dinoflagellates and horizontally-transfer algae. Molecular characterizations of two such anthozoans, a soft coral *Xenia* species and a hard coral *Stylophora pistillata*, revealed that the alga-host cells express specific lectins, scavenger receptors, and phagocytosis machinery for algal uptake[6, 7]. It remains unknown, however, whether algal uptake in cnidarians performing facultative endosymbiosis exhibit the same features. Interestingly, some cnidarians, including the reef-building corals and *Hydra viridissima* (*H. viridissima*), also evolved the ability to take up algae during oogenesis, which endows the next generation with symbionts[8, 9]. The benefits of this vertical algal transmission remain unclear. Since *H. viridissima* performs facultative endosymbiosis and vertical algal transmission, we have used genomics and cell biology approaches to uncover the regulatory features of this poorly understood evolutionary adaptation.

## Results

### Cell-type classification aided by single cell RNA-sequencing

We performed single-cell RNA-sequencing (scRNA-seq) of enzymatically-dissociated[10, 11] *H. viridissima* with or without developing egg patches. The *Hydra* dissociation procedure yielded a mix of free algae, small (∼6-7 μm) and large *Hydra* cells (15-20 μm) (Figures S1A and S1B), with some large intact cells, as assessed by surface biotinylation and Streptavidin labeling, containing 1 to 37 algae[12, 13] (Figures S1C and S1D). Cell clustering of the scRNA-seq data identified 32 clusters (Figures 1A-1C). To identify *H. viridissima* cell types, we performed a cross-species comparison with a previous scRNA-seq annotation for *Hydra vulgaris* (*H. vulgaris*)[14], a non-endosymbiotic relative of *H. viridissima* (Figures 1D and S1E). The <u>En</u>doderm <u>Ep</u>ithelial <u>S</u>tem <u>C</u>ell (enEpSC; clusters 0, 2, 3) and <u>Ec</u>toderm <u>Ep</u>ithelial <u>S</u>tem <u>C</u>ell (ecEpSC; clusters 1, 4)[14] represent the major cell clusters in *H. viridissima* (Figures 1A and 1B). We also identified several neuronal cell clusters (13, 17, 21, 26, 30), two nurse cell clusters (7, 14), primarily in *H. viridissima* bearing gonads, and the interstitial <u>s</u>tem <u>c</u>ells (iSC, cluster 5) (Figures 1A-1D). The *H. viridissima* iSC cluster shows the strongest similarity to the *H. vulgaris* iSC followed by neuron and gland progenitors, which are both derived from the iSC (Figures 1D and S1E)[14, 15]. We next performed Fluorescence *In Situ* Hybridization (FISH) using probes against the gene (scaffold35.g35) that is specifically expressed in the nematoblast cell cluster 16 (Figure S1F) and confirmed its specific expressed in clusters of cells in the ectoderm where nematoblasts reside (Figure 1E).

**Figure 1.**
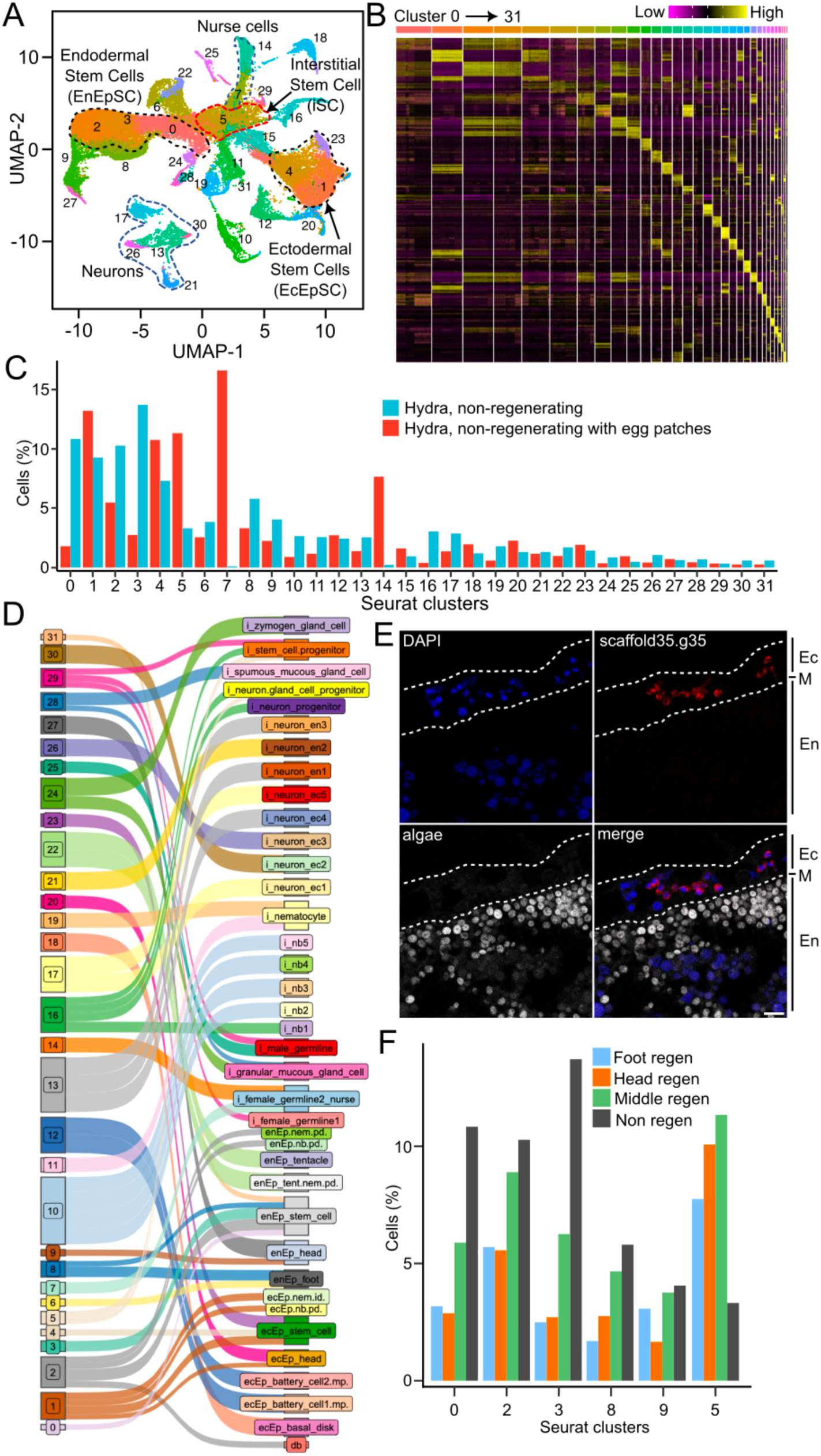
Cell type classification in *H. viridissima*. **A**. Uniform Manifold Approximation and Projection (UMAP) plot of 32 cell clusters (0 to 31) identified from scRNA-seq of *H. viridissima*. **B.** Heatmap showing expression patterns of marker genes across the 32 (0 to 31) identified Seurat clusters. **C.** Bar-chart showing the percentage of cells in each cluster from whole *H. viridissima* with (red) or without egg patches (blue). **D.** Sankey diagram illustrating the correspondence between *H. viridissima* cell clusters (left) and annotated cell types from *H. vulgaris* (right). **E.** Fluorescence *in situ* hybridization (FISH) validating the specific expression of scaffold35.g35 in nematoblast cells (cluster 16). DAPI (blue) stains nuclei, scaffold35.g35 probe (red) shows expression in a linear string of cells in the ectoderm (Ec) as outlined by dotted lines. Algal autofluorescence (grey) is visible in the endoderm (En). M indicates mesoglea. **F.** Bar-chart showing the percentage of cells in selected clusters (0, 2, 3, 8, 9, 5) in non-regenerating animals (“Non regen”, black) and animals undergoing foot regeneration (“Foot regen”, blue), head regeneration (“Head regen”, orange), or middle body regeneration (“Middle regen”, green). Note the reduction of enEpSC clusters 0, 2, and 3 and the increase in iSC cluster 5 in regenerating samples.

Studies of *H. vulgaris* suggest that enEpSC and ecEpSC primarily support the differentiation and maintenance of their respective tissue layers, while the iSCs differentiate into the secretory precursor cells, nematoblasts, gland and nerve cells[15–20]. *H. vulgaris* regeneration may rely on the tissue-specific enEpSC and ecEpSC, but not the iSC[15, 19]. Our scRNA-seq on *H. viridissima* undergoing head, foot, or both the head and foot (middle) regeneration showed, however, that animals undergoing regeneration had a striking reduction of enEpSC clusters 0, 2, and 3, whereas the number of cluster 5 iSCs showed a >2-fold increase in regenerating animals compared to non-regenerating ones (Figure 1F). The increase in the iSCs during regeneration in *H. viridissima* may replenish the lost enEpSCs to aid regeneration. In support of this, we analyzed the lineage differentiation trajectory with Slingshot using the cluster 5 iSCs as the differentiation start point followed by the enEpSC clusters 0, 2 and 3[21]. Indeed, the iSCs could give rise to enEpSCs cluster 0 followed by 3 and then 2 (Figure S1G).

### Identification of three endoderm cell types as the major hosts for *Chlorella* algae

We used Fluorescence Activated Cell Sorting (FACS, Figure 2A, left) to enrich for intact, putative algae-containing cells (Figure 2A, right), based on their strong algal autofluorescence, for bulk RNA-seq. Compared with individual scRNA-seq cell clusters, we unexpectedly found that many cell types may contain algae (Figure 2B). We then performed scRNA-seq of these FACS-isolated cells and mapped them to the scRNA-seq dataset. Again, we found a similar set of putative algal containing cells with the strongest representation in endoderm clusters 0, 2, 3, 6, 8, 9 and ectoderm cluster 15 (Figures 2C and S2A). Using Ce3D tissue clearing (see Methods) and an anti-Calnexin antibody that strongly labels the mesoglea and weakly labels all cell boundaries (Figure 2D), we found most algae resided within irregularly-shaped columnar endodermal cells under the mesoglea (Figure 2D, yellow dashed outlines). This is consistent with previous reports.[5, 22, 23]. We also found free algae, albeit rarely, in the gastrodermis cavity (Figure 2D, white dashed outline)[24, 25]. Since imaging of dissociated Hydra and FACS-isolated putative alga-host cells showed that algae could be attached to the surface of some algal-free cells (Figure S1A, yellow arrows), individual droplets in the subsequent scRNA-seq procedure could contain both animal cells and free algae and result in the misidentification of some algal host cells.

**Figure 2.**
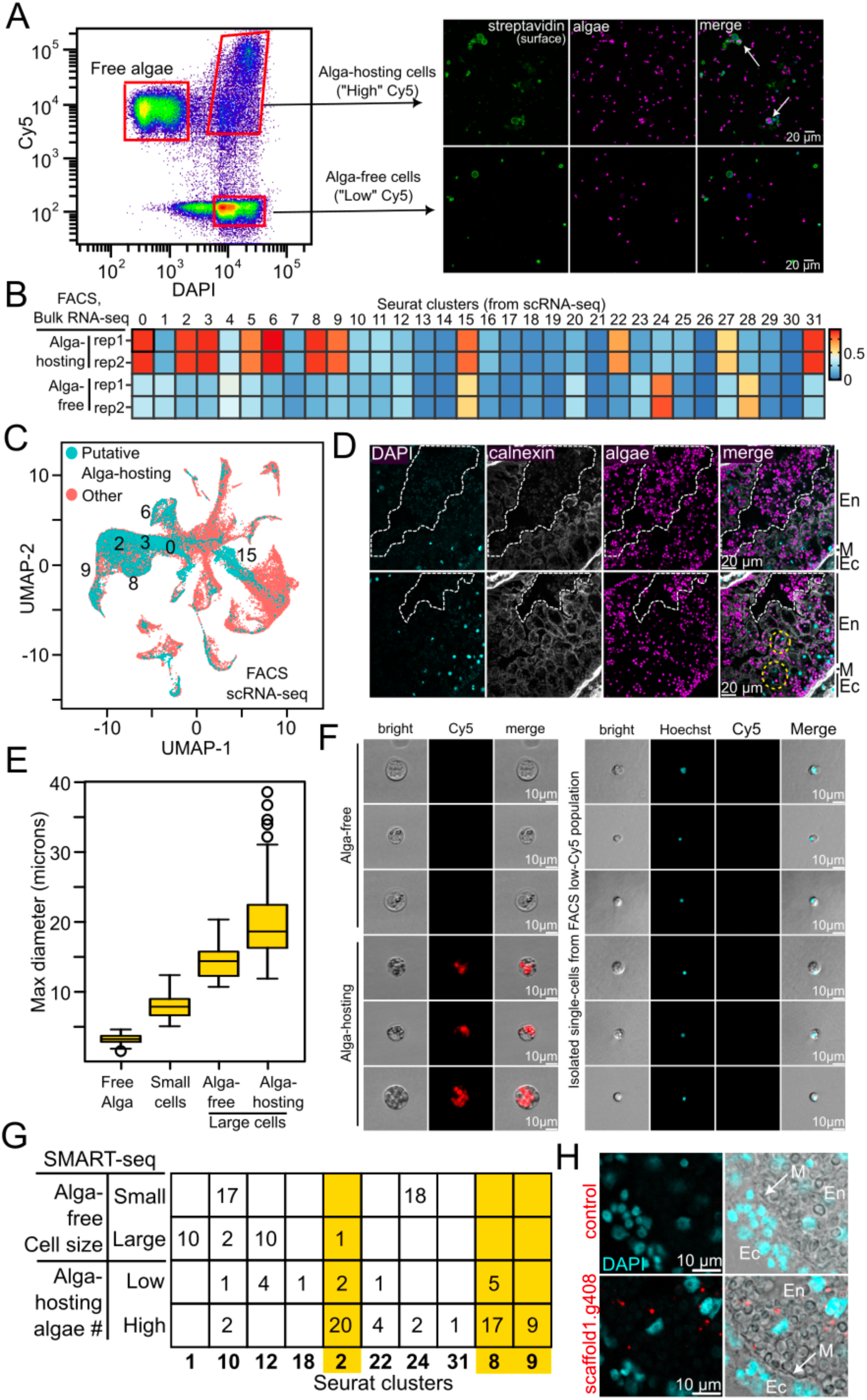
Identification of algal-hosting cells in *H. viridissima*. **A.** FACS plot (left panel) showing the gating strategy for isolating alga-hosting cells (“High” Cy5) and alga-free (“Low” Cy5) animal cells from dissociated *H. viridissima*. DAPI indicates DNA content. Right panel showing FACS-isolated cells that were surface biotinylated with EZ-link sulfo-NHS-biotin and stained with DAPI and Streptavidin (green). Algal autofluorescence is in magenta. Arrows point to examples of alga-hosting cells isolated by the FACS gating strategy. Scale bar = 20 µm. **B.** Heatmap showing the correlation between bulk RNA-seq profiles from FACS-isolated *H. viridissima* cells having (“alga-hosting”) or not having algal autofluorescence (“algal-free“) and the transcriptomes of individual scRNA-seq cell clusters. Clusters 0, 2, 3, 5, 6, 8, 9, 15, 22, 27, and 31 show high correlation with the putative algal containing cells. **C.** UMAP plot showing the distribution of scRNA-seq of FACS-isolated putative alga-hosting cells (cyan color) overlaid on the UMAP of all cell clusters (red) from the overall scRNA-seq dataset. Matching clusters are labeled. **D.** Confocal microscopy images of whole-mount Ce3D cleared *H. viridissima* stained with DAPI (nuclei, cyan), anti-Calnexin antibody (labelling mesoglea and cell boundaries, greys) and algal autofluorescence in magenta. Images show algae primarily residing within large endodermal cells (dashed yellow outlines). Some free algae appear in the gastrodermal cavity (outlined in dashed-white). Black bars on the outside of the image mark the approximate tissue regions. “Ec” indicates ectoderm, “M” mesoglea and “En” the endoderm. Scale bar = 20 µm. **E.** Boxplot showing the maximum diameter (microns) of free algae, small alga-free cells (Small cells), large alga-hosting cells (Alga-hosting/Large cells), and large, spherical alga-free cells (Alga-free/Large cells). **F.** Images of hand-isolated single cells used in the SMART-seq experiment. Left panel shows large alga-free (“Alga-free”) and alga-hosting (“Algal-hosting”) cells. Bright-field is in greys and algal autofluorescence (Cy5, red). The right panel shows small cells isolated after FACS sorting that also include Hoechst staining of nuclei (cyan). Scale bar = 10 µm. **G.** Table summarizing SMART-seq results of hand-picked small and large alga-free cells, and verified algal-hosting cells containing high (>5 algae) or low (<5 algae), mapped to scRNA-seq clusters. The major large alga-containing cells in Clusters 2, 8 and 9 are highlighted in orange. **H.** FISH showing the scaffold1.g408 (red) positive cells containing algae as indicated by their appearance in the bright-field channel. DAPI (cyan) stained for nuclei. White arrow, mesoglea (M). Scale bar = 10 µm.

To definitively identify the algal containing cells, we analyzed the size of dissociated individual *H. viridissima* cells and found that small and large cells could be broadly distinguished (Figure S1A and S1B). Dissociated algae-free cells could be small (∼6-7 μm) or large (∼15 μm), and all algae-containing cells are large (>17 μm) in diameter (Figure 2E). We next hand-picked large cells with or without algae (Figure 2F, left panels) or small cells (Figure 2F, right panels) and performed SMART-seq[26]. Since cell debris and free algae make it difficult to isolate true small cells, we first used FACS (Figure 2A) to gate for small cells and then hand-picked cells with interior features. Mapping the SMART-seq transcriptome to the scRNA-seq above identified the enEpSC cluster 2, as the most abundant algal containing cells followed by endoderm cell clusters 8 and 9 in the foot and head, respectively (Figure 2G). FISH analyses with a gene (scaffold1.g408), most strongly expressed in the algal containing enEpSC cluster 2 cells (Figure S2B), showed that these cells are in the endoderm (Figure 2H and S2C).

Our SMART-seq identified cells in clusters 1 as non-algal containing large cells, cluster 10 and 24 as mainly small non-algal containing cells, and cluster 12 as large cells with no or small numbers of algae (Figure 2G). The SMART-seq did not identify any cells from some of the most abundant cell clusters 0, 3, 4, and 5 (Figure 1A). This could be due to technical issues related to cell isolation or limited sampling. Interestingly, clusters 10 (nematoblasts), 12 (battery cells), and 18 (ecEp basal disk) are ectoderm cells, but our SMART-seq showed that some of these cells contain a few algae (Figure 2G). Consistently, we found that some ectoderm cells in different regions of *H. viridissima* contained one and often misshapen alga based on electron microscopy (Figure S2D)[27]. Since individual *Chlorella* algae found in the endoderm often appear round and healthy (Figure S2B), these ectoderm cells may occasionally take up algae via constitutive particle uptake pathways all cells have, but may not be able to maintain them[28]. Thus, the definitive endosymbiotic cells in *H. viridissima* are the enEpSC cluster 2 and endoderm cell clusters 8 and 9.

### Molecular signatures of algal uptake and maintenance in the algal host cells

Unlike the findings in the soft and hard corals performing obligate endosymbiosis[6, 7], we found no marker genes encoding lectin, scavenger receptor and algal uptake proteins in *H. viridissima* algal host cells that could explain a specific machinery for establishing endosymbiosis[6, 7, 29, 30]. To support the facultative symbiosis lifestyle, however, *H. viridissima* needs to take up algae. We explored whether constitutive phagocytic or endocytic activities in *Hydra* cells may enable algal uptake and found that the endosymbiotic cell clusters 2, 8, and 9 are among those having the highest receptor-mediated endocytic and phagocytic activities (Figures 3A and 3B). Thus, constitutive uptake activities may be one mode used to take up algae, possibly by many cells, but only the endosymbiotic cells can maintain algae. We did, however, find that in the apo-symbiotic state the algal host cells express specific genes (e.g., lectins) that may be involved in efficient algal uptake (data not shown), which suggests at least two modes of uptake depending on symbiotic or aposymbiotic state.

**Figure 3.**
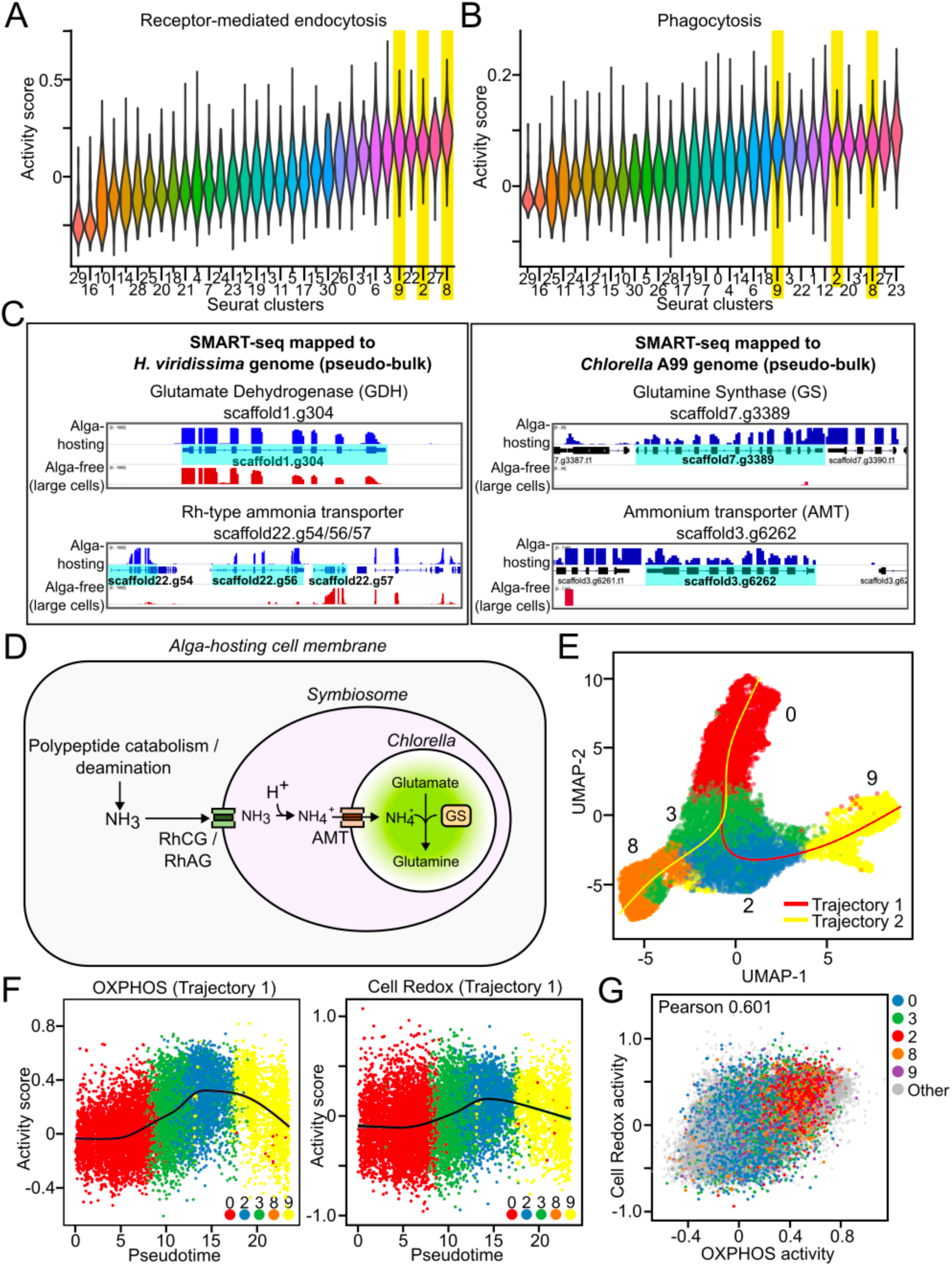
Molecular signatures of algal-hosting cells. **A., B.** Violin plots illustrating the activity score for receptor-mediated endocytosis (**A**) and phagocytic (**B**) in each cell cluster. Clusters 2, 8, and 9 are among those with the highest activity and highlighted in yellow. **C.** IGV genome browser tracks showing expression levels of *H. viridissima* genes encoding (left panels) glutamate dehydrogenase (GDH, scaffold1.g304) and Rh-type ammonia transporters (scaffold22.g54, 56, 57). The *Chlorella* genes (right panels) encoding glutamine synthase (GS, scaffold7.g3389) and ammonium transporter (AMT, scaffold3.g6262) in the large alga-hosting and alga-free cells from the SMART-seq data. These data are presented as a pseudo-bulk of SMART-seq transcriptomes. The gene model is highlighted in cyan. RNA-seq data for large alga-hosting cells (or algae inside) and alga-free cells is colorized in blue and red, respectively. **D.** Model for ammonia provision from algal host cell to algae. Host cell polypeptide catabolism and deamination produce ammonia. Host cell Rh-type ammonia transporters (e.g., RhCG/RhAG homologs) transport ammonia (NH_3_) across the symbiosome membrane into the acidic symbiosome where it becomes ammonium (NH_4_). Algal ammonium transporter (AMT) takes up ammonium, which is then assimilated into glutamine by algal glutamine synthase (GS). **E.** UMAP plot showing Slingshot differentiation trajectories of algae-hosting cells and their progenitors. Two trajectories leading to cluster 8 or 9 are labeled. **F.** Scatterplot showing the OXPHOS activity (left panel) and redox homeostasis activity (right panel) along the alga-hosting cell maturation trajectory 1. **G.** Scatterplot showing the correlation (Pearson 0.601) between OXPHOS activity and redox homeostasis activity across all cells. Most alga-hosting cells (cluster 2, 8, 9) show high OXPHOS and redox activity.

The *Chlorella* symbionts in *H. viridissima* have lost several genes involved in nitrogen assimilation[31]. Studies based on microarray of symbiotic and aposymbiotic *H. viridissima* suggest that *Hydra* cells could supply glutamine to algal endosymbionts[31]. We found that endosymbiotic cells 2, 8, and 9 do not show a specific high glutamine biosynthesis activity (Figure S3A), but they exhibit an increased protein degradation activity (Figure S3B and S3C). SMART-seq provided transcriptomes for individual *Hydra* cells and algae inside *Hydra* cells (Figure S3D). Further analyses showed that the large algal host and algal free cells both express genes such as Glutamate Dehydrogenase (GDH), but algal host cells specifically express two of three Rh-type ammonia transporters clustered on scaffold22 (scaffold22.g54, scaffold22.g56) (Figure 3C, S3D-F). Importantly, the *Chlorella* symbionts in the algal host cells express both ammonium transporter (AMT, scaffold3.g6262) and Glutamine Synthase (GS, scaffold7.g3389, Figure 3C, S3D)[32]. Interestingly, the expression of the two Rh-type ammonia transporters was reduced in apo-symbiotic *H. viridissima* algal host cells compared to those in the symbiotic animals (data not shown). Ammonia transported into the symbiosome of host cells could be converted into ammonium in the acidic environment of the symbiosome, which can then be transported into alga by algal AMT and used to make glutamine by algal Glutamine Synthase (Figure 3D).

### Strong signatures for ATP production by oxidative phosphorylation and redox in algal host cells

We analyzed the Slingshot lineage differentiation trajectory[21] with enEpSC clusters 0, 2, 3, and endoderm cell clusters 8 and 9. The enEpSCs cluster 0 gives rise first to 3 and then 2. Cells in clusters 3 and 2 then give rise to the foot and head cell clusters 8 and 9, respectively (Figure 3E). Using cell activity score analyses (see Methods)[33, 34], we found a gradual increase in activities of ATP production by oxidative phosphorylation (OXPHOS) and redox along the lineage trajectory (Figure 3F trajectory 1, S4A, and S4B trajectory 2). The algal containing cluster 2 cells exhibited the highest scores followed by some cells in cluster 8 and 9. While the two activities had a Pearson correlation of 0.601 in all cells, the algal containing cluster 2 cells exhibited the highest activities followed by some cells in cluster 8 and 9 (Figure 3G).

The increased OXPHOS-ATP production in the mitochondria by the endosymbiotic cells could be supported by various carbon compounds, including sugar, produced by algal photosynthesis. The Reactive Oxygen Species (ROS) generated by OXPHOS in endosymbiotic host cells could in turn elevate a redox transcriptional response to preserve cellular and mitochondria function[35, 36]. There is a reduced OXPHOS and redox activities toward the end of the lineage trajectory, which could be caused by the persistent OXPHOS and increased ROS that eventually overwhelms the redox system. The transcriptional responses in OXPHOS and redox by the algal host cells have not been previously reported in animal-algal endosymbiotic cells and it may reflect an evolutionary adaptation that enables *H. viridissima* to derive carbon sources from algal photosynthesis to produce energy in the form of ATP. Interestingly, some studies showed that the aposymbiotic *H. viridissima* had reduced egg production[13]. The OXPHOS ATP production by algal containing cells may have been evolved in part to support the energy demanding oogenesis in *H. viridissima*.

### Cellular features of vertical algal transmission

We found that the nurse cell clusters express a plant-derived peroxidase gene (scaffold59.g26, “hvAPX”) as reported previously[37] (Figure S5Aa). Using the peroxidase activity and algal autofluorescence, we found an alga increase in the body column under the egg patches starting at stage 2 (total 7 stages)[38–40] of oogenesis, which persisted until egg hatching (Figure 4A). These changes may support algal uptake by the developing oocytes for vertical algal transmission. The earliest time algae appear in the oocytes is stage 3 (Figure S5B; Video S1) but is more evident and numerous at stage 4 and beyond (Figure 4B, white arrows).

**Figure 4.**
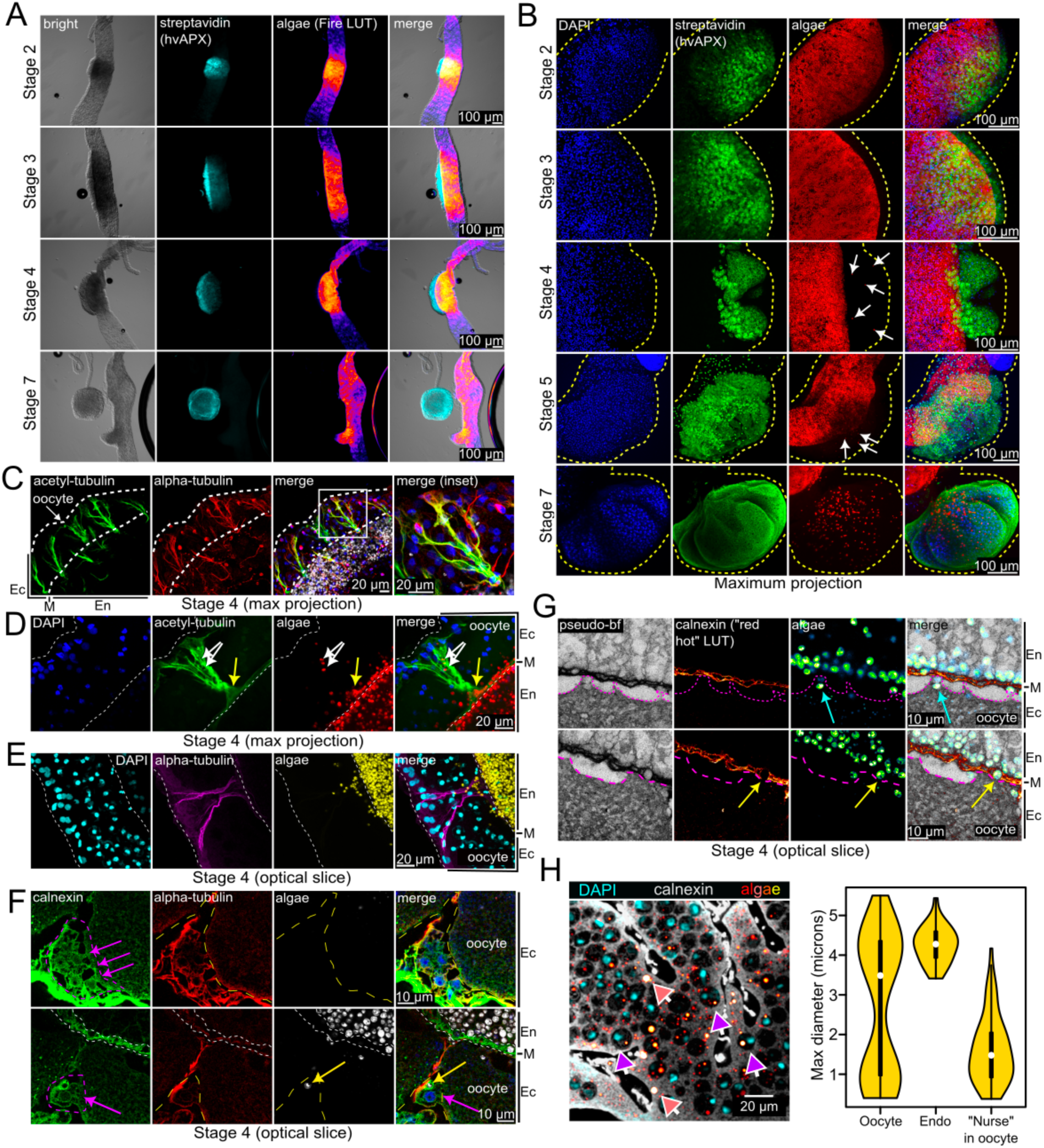
Cellular features of vertical algal transmission. **A.** Visualization of nurse cells by streptavidin staining after the biotin-phenol peroxidase reaction (“streptavidin, hvAPX”, cyan) and algal autofluorescence (signal intensity shown with Fire LUT) in *H. viridissima* during stages 2-7 of oogenesis. An increase in algae is observed in the body column under the egg patch from stage 2 onward. Bright-field is shown in greys. Scale bar = 100 µm. **B.** Algae appear in the oocyte at stage 3 and are more numerous at stage 4. DAPI indicates nuclei, Streptavidin (“hvAPX”, green), and algae (red). White arrows point to algae visible within the space occupied by the developing oocyte. Scale bar = 100 µm. **C.** Long cell processes with microtubules appear to originate from the surface of the developing egg patch and end at the mesoglea in a stage 4 egg patch. Acetylated tubulin is in green and alpha tubulin in red. The merged panels contain DAPI staining (blue) and algae (greys) not presented as separate panels. The white box highlights the zoomed inset on the right. The ectoderm (Ec) and endoderm (En) are marked by black-bars on the outside of the first panel. White dashed lines enclose the egg patch. The image is a max projection and scale bar = 20 µm. **D.** Multiple microtubule processes converging into a bouquet-like structure in a stage 4 egg patch as illustrated by max projection. DAPI is presented in blue, acetylated tubulin in green and algae in red. White arrows point to algae within microtubule bouquet and the yellow arrow points to a cluster of algae at the base of the bouquet. Scale bar = 20 µm. **E.** An optical slice of a stage 4 oocyte showing algae along the microtubule track. DAPI is in cyan, alpha tubulin is in yellow and algae in yellow. Scale bar = 20 µm. **F.** Two optical confocal slices of an egg patch. White dashed lines demarcate mesoglea. The top panel is a deeper plane showing connections between a cluster of putative small nurse cells (purple dashed outline) connected to the oocyte (purple arrows). The bottom panel is from an upper plane showing the position of an algae (yellow arrow) on the microtubule track (red) juxtaposed to a putative nurse cell (purple dashed outline). The putative nurse cell appears to have extensions that are in contact with the oocyte (purple arrows). DAPI (blue), Calnexin (green), Alpha-tubulin (red) and algae (greyscale). Scale bar = 10 µm. **G.** Confocal microscopy images of a stage 4 developing oocyte. The pseudo-bright field (“pseudo-BF”) is a grayscale image of the Calnexin stain at 488nm and the autofluorescence in the 568nm channel that have been contrasted. Calnexin staining, non-contrasted, is presented in “red hot LUT” to highlight the mesoglea (M) and free algae are as a green LUT. Free algae are visible within the mesoglea and juxtaposed to it on the ectodermal (Ec) side of the oocyte. Yellow arrows indicate an apparent break in the Calnexin-demarcated mesoglea where an alga appears to have entered. The cyan arrows point to two algae that appear to be inside an oocyte-derived membrane protrusion. Oocyte-derived membrane protrusions are outlined by dashed magenta lines, and seen contacting the mesoglea. Scale bar = 10 µm. **H.** Confocal image of an area of a stage 5 oocyte (left panel) stained with DAPI (nuclei, cyan), Calnexin (cell boundaries, greyscale), and algal autofluorescence (red hot LUT). Many DAPI-positive spherical bodies, identified as engulfed nurse cells, are visible within the oocyte. White-bordered pink arrows point to large algae in the presumed oocyte cytoplasm. White-bordered purple arrows point to presumed nurse cell bodies containing DAPI-positive nuclei and algae autofluorescence signal. Scale bar = 20 µm. Violin plot (right panel) showing the sizes of algae in the oocyte cytoplasm (Oocyte), endoderm algal host cells (Endo), and in nurse cells (“Nurse” in oocyte). A population of algae in the oocyte cytoplasm are similar in size to those seen in the algal host cells. Ectoderm (Ec), Mesoglea (M) and Endoderm (En) are marked by black-bars on the outside of relevant images.

It was proposed that algae could be delivered to the oocyte by cell processes originating from endoderm cells or by passive diffusion[8, 41]. However, using antibodies to tubulin and acetylated tubulin, we found long cell processes that appeared to originate from the surface of developing egg patch and ended at the mesoglea (Figure 4C, Videos S2 and S3). Multiple cell processes with microtubules could converge into what appeared as a “bouquet-like” structure (Figure 4D, Video S4) with algae often found at the mesogleal interface of this structure and along microtubule processes (Figure 4D and 4E). We did, albeit rarely, observe algae (yellow arrows) on the microtubule process near what appeared to be nurse cells (outlined by purple dashed lines) attached to the oocyte (purple arrows)[38, 42] (Figure 4F, Video S4). Labeling of cell boundaries and mesoglea with the anti-Calnexin antibody combined with Ce3D tissue clearing (Figure S5C and Video S5) revealed free algae within the mesoglea layer and also juxtaposed to it on the ectodermal side of the developing oocyte (Figure 4G and Video S6). Some Calnexin demarcated mesoglea appeared broken in regions where algae had entered the mesoglea (Figure 4G, yellow arrows). We also observed that the developing oocyte frequently contacts the mesoglea via membrane protrusions, and some membrane protrusions have an alga inside or contact alga in the broken mesoglea (Figure 4G, blue arrows and Video S6). Inside the stage 4-5 oocyte, we found many spherical bodies with each containing a nucleus as based on DAPI staining (Figure 4H left panel, purple arrows). These DAPI-positive bodies are nurse cells engulfed by the oocytes[40, 42]. The nurse cell bodies often contain 1 or more algae as judged by autofluorescence. The algae are often smaller (1.6 μm +/-0.83) than those found in the endoderm algal host cells (4.2 μm +/- 0.45), while the algae in the cytoplasm of developing oocytes (Figure 4H left panel, pink arrows) are either small (0.9 μm +/- 0.29) as seen in the nurse cells or large as seen in the algal host endoderm cells (4.2 μm +/- 0.53) (Figure 4H, right panel). Interestingly, the small-sized algae in the oocytes and the nurse cell bodies are about half in size of the large algae in the oocytes. The small algae in nurse cells may be in G1 phase and upon release into the oocyte cytoplasm, they may progress into G2 phase and proliferate. Alternatively, the smaller algae in the internalized nurse cell may be undergoing degradation and only some algae taken up by the oocytes may be transferred to the next generation. Thus, the microtubule processes could be involved in transferring algae from the endoderm to the developing egg patch to be taken up by the oocyte and nurse cells.

### Molecular characteristics of cells and genes supporting oogenesis

We next performed scRNA-seq by dissecting out the body column bearing stage 1-3 or stage 4-5 oocytes (Figure 5A and 5B). Comparing to the full animal scRNA-seq, we found an increase in nurse cells in clusters 7 and 14, and a reduction of algal host cells in clusters 9 and 8 in these samples, as expected (Figure 5C). Interestingly, the egg patch samples had an enrichment of cluster 0 cells and a decrease in cluster 2 algal-containing cells (Figure 5C). These cluster 0 cells have elevated activity scores for extracellular matrix (ECM) breakdown, especially in the stage 4-5 oocyte samples when most of cluster 0 cells express these genes (Figure 5D and 5E). ECM breakdown could aid algae transfer from the endoderm to the egg patch as observed (Figure 4G). The cluster 2 algal host cells may support vertical transmission by releasing algae and become depleted afterward (Figure 5C). We further analyzed scRNA-seq of stage 1-3 and stage 4-5 egg patch fragments and found that both nurse cells and many other cell types upregulate apolipoprotein B (APOB) (Figure 5F). APOB may be involved in transporting lipid into *H. viridissima* oocytes during oogenesis. Interestingly, in *Drosophila* and mice, APOB derives lipids from organs such as fat body (*Drosophila*) or liver (mice), which are then supplied to their developing oocytes[43–46]. The greatly simplified body parts in *H. viridissima* may have driven the evolution of APOB expression in many cell types during oogenesis to supply the oocytes with needed lipid storage.

**Figure 5.**
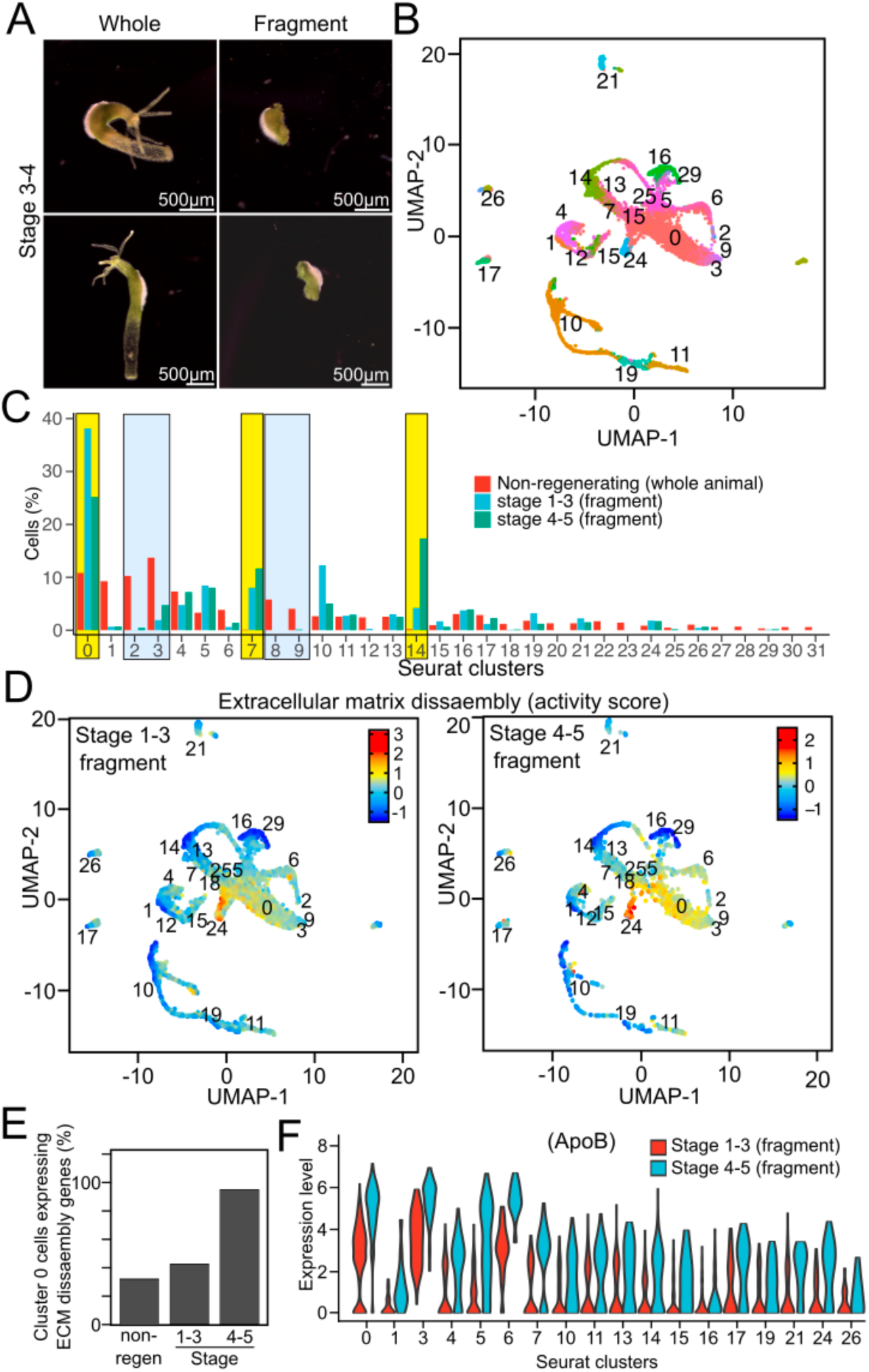
Cells and genes supporting oogenesis and algal delivery. **A.** Examples of *H. viridissima* bearing stage 3-4 egg patches (Whole) and dissected body column containing the egg patch (Fragment). Scale bar = 500 µm. **B.** UMAP clusters for the integrated oogenesis scRNA-seq datasets for stage 1-3 and stage 4-5. The cell cluster numbers correspond to the full body scRNA-seq. **C.** Bar-chart comparing cell percentages observed for whole, non-regenerating (“Non-regenerating (whole animal)”) and the oocyte bearing fragments of *Hydra* at the indicated stages, stage 1-3 (fragment) and stage 4-5 (fragment). Clusters 0, 7 and 14 were enriched in the egg-patch fragment samples and are highlighted in yellow. Clusters 2, 3, 8 and 9 were decreased in egg-patch samples and are highlighted in blue. **D.** UMAP with heat map showing the activity score for extracellular matrix disassembly in the Stage 1-3 and Stage 4-5 fragments scRNA-seq. **E.** Bar-chart showing the percentage of cells expressing Extracellular matrix disassembly (“ECM”) genes in cluster 0 of non-regenerating animals (“non-regen”), stage 1-3 and stage 4-5 scRNA-seq. **F.** Many cells, including the nurse cells in clusters 7 and 14, exhibit upregulation of apolipoprotein B (ApoB, scaffold58.g58) at stage 4-5 compared to the Stage 1-3 fragment.

## Discussion

Here we identify the cell types in *H. viridissima*, including three populations of endosymbiotic cells. We found that the algal-hosting cells in the symbiotic *H. viridissima* do not specifically express genes involved in algal recognition and uptake, but they have high constitutive phagocytic and endocytic activities, which may support algal uptake. We also identify specific expression of Rh-type ammonia transporter genes in the algal host cells that can enable the algae to derive ammonium from the host cells. There is an increase in algae in the body column under the developing oocyte and algal entry into the stage 3-4 oocyte. Since algal number continue to increase in the developing oocytes, vertical algal transmission may have evolved to leverage photosynthesis to support the energy demands of sexual reproduction and embryogenesis. Consistently, studies show that the aposymbiotic *H. viridissima* have reduced egg patch formation[13]. Additionally, the increased OXPHOS ATP production and redox activity by the algal host cells may enable *H. viridissima* to adapt to a nutrient poor niche during its evolution. Together, our findings should enable molecular studies of facultative endosymbiosis and vertical transmission in *H. viridissima*. Indeed, RNAi tools have been developed to study *H. vulgaris* that does not perform endosymbiosis but share a lot of similarities with *H. viridissima*, including the RNAi machinery. To study gene function in the algal containing cells, it is important to use cell-type specific gene knockdown. Our scRNA-seq and SMART-seq should facilitate the identification of tissue specific promoters, which would enable cell-type specific gene silencing.

## Acknowledgements

We thank Dr. Frederick. Tan (Carnegie / Johns Hopkins University), Dr. Javier Carpinteyro-Ponce (Carnegie) and Allison Pinder (Carnegie) for assistance with sequencing and initial data processing; Lynne Hugendubler (Carnegie) for maintaining the hydra culture; Dr. Asya Davidian (Carnegie) for suggestions on imaging and tissue clearing; Dr. Mahmud Siddiqi (Carnegie) for help with FACS, confocal and electron microscopy; Dr. Xin Chen (Johns Hopkins University) for use of the individual cell-isolation rig, and Drs. Sveta Deryusheva and Alex Bortvin (Carnegie) for creating and providing the apo-symbiotic *H. viridissima*.

## Author contributions

J.R.T., J.S., M.H. and Y.Z. conceived the project. J.R.T., J.S., M.H., and Y.Z. designed experiments. J.R.T., J.S., and M.H. performed 10x scRNA-seq (M.H., J.R.T., J.S.), bulk RNA-seq (J.S.), and SMART-seq (J.R.T. and B.M.) experiments. J.S. performed initial scRNA-seq and bulk RNA-seq analysis. M.H. performed additional scRNA-seq experiments using FACS material and regenerating *Hydra* and data analyses. J.R.T. performed the SMART-seq data analyses and the immunofluorescence and imaging. M.H. performed the electron microscopy. M.Z. performed the FISH experiments. J.S., J.R.T., M.H, and Y.Z. analyzed the data. J.S., J.R.T., M.H., and Y.Z. interpreted the data. J.R.T. and M.H. prepared the figures, movies and supplemental tables. Y.Z. wrote the initial draft. J.R.T., M.H., and Y.Z. edited the draft. All authors reviewed and edited the manuscript.

---

**Figure S1.**
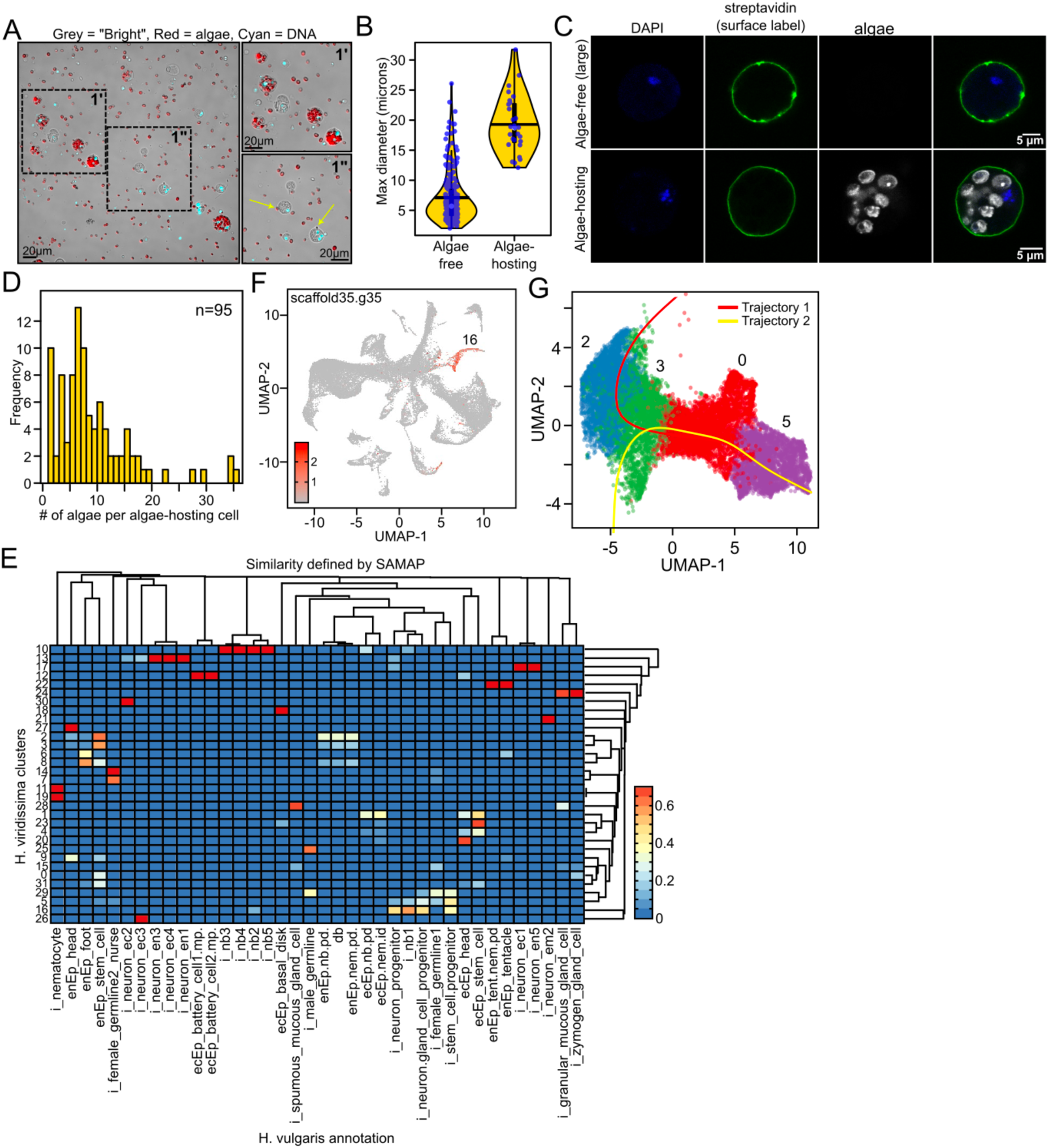
Additional data for *H. viridissima* scRNA-seq. **A.** Representative images of dissociated *H. viridissima* cells used for scRNA-seq. Insets (1’ and 1”) are denoted by dashed-line squares. Yellow arrows point to non-algal host cells associated with alga on the surface. Bright-field is in grayscale, DAPI in cyan and algal autofluorescence in red. Scale bar = 20 µm. **B.** Violin plot of cell size distribution. **C.** The cell surface is intact in dissociated Hydra cells. The cell surface was labeled with EZ-link sulfo-NHS-biotin and detected with Streptavidin (green). DAPI is in blue and algae autofluorescence in greyscale. Scale bar = 5 µm. **D.** Histogram plot showing the distribution of algae number in individual symbiotic cells from the dissociation procedure. **E.** SAMAP heatmap comparing *H. viridissima* cell clusters with annotated cell types from *H. vulgaris*. **F.** UMAP plot showing the expression of a nematoblast marker gene (scaffold35.g35) in cluster 16. **G.** UMAP plot showing Slingshot differentiation trajectories (Trajectory 1 and 2) among iSC (cluster 5) and enEpSC clusters (0, 3, 2).

**Figure S2.**
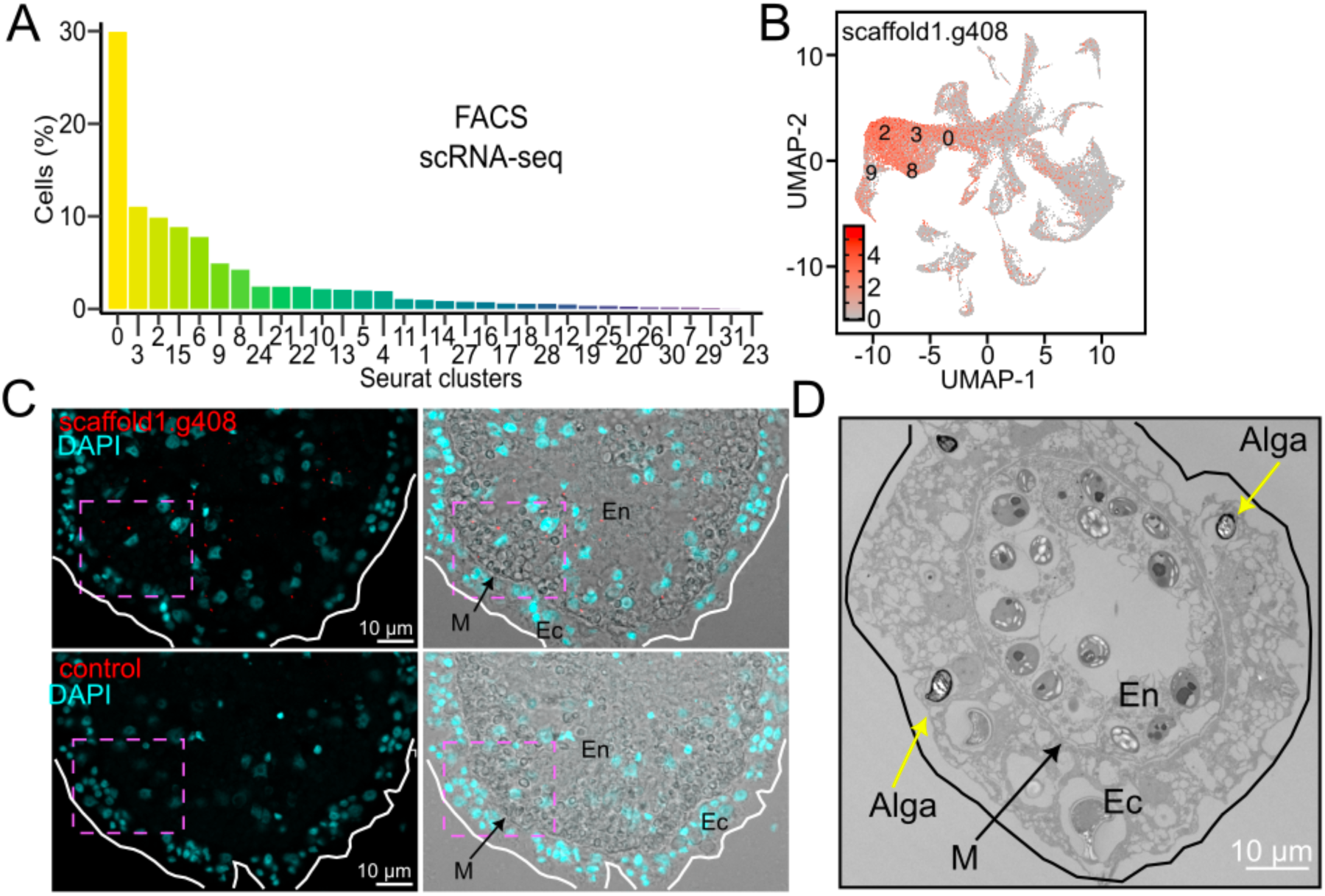
scRNA-seq of FACS isolated cells and tissue sections. **A.** Bar-chart showing the cell percentage of different Seurat clusters identified from scRNA-seq of FACS-isolated cells with high-algal autofluorescence. Clusters 0, 2, 3, 6, 8, 9, and 15 are best represented. **B.** UMAP plot showing the expression of scaffold1.g408, a marker gene most enriched in the labeled endoderm cells with highest enrichment in cluster 2 cells. **C.** FISH image showing the localization of a gene (scaffold1.g408) in red. Cells containing algae can be seen in the bright-field channel. DAPI (cyan) stained for nuclei. Insets used in Figure 2g are denoted with a pink dashed-line square. Scale bar = 10 µm. **D.** Electron microscopy image showing algae within ectoderm (Ec) and endoderm (En) cells. The black arrow points to the mesoglea (M). Yellow arrows point to algae in ectoderm that appear smaller or misshapen compared to those in the endoderm cells. Scale bar = 10 µm.

**Figure S3.**
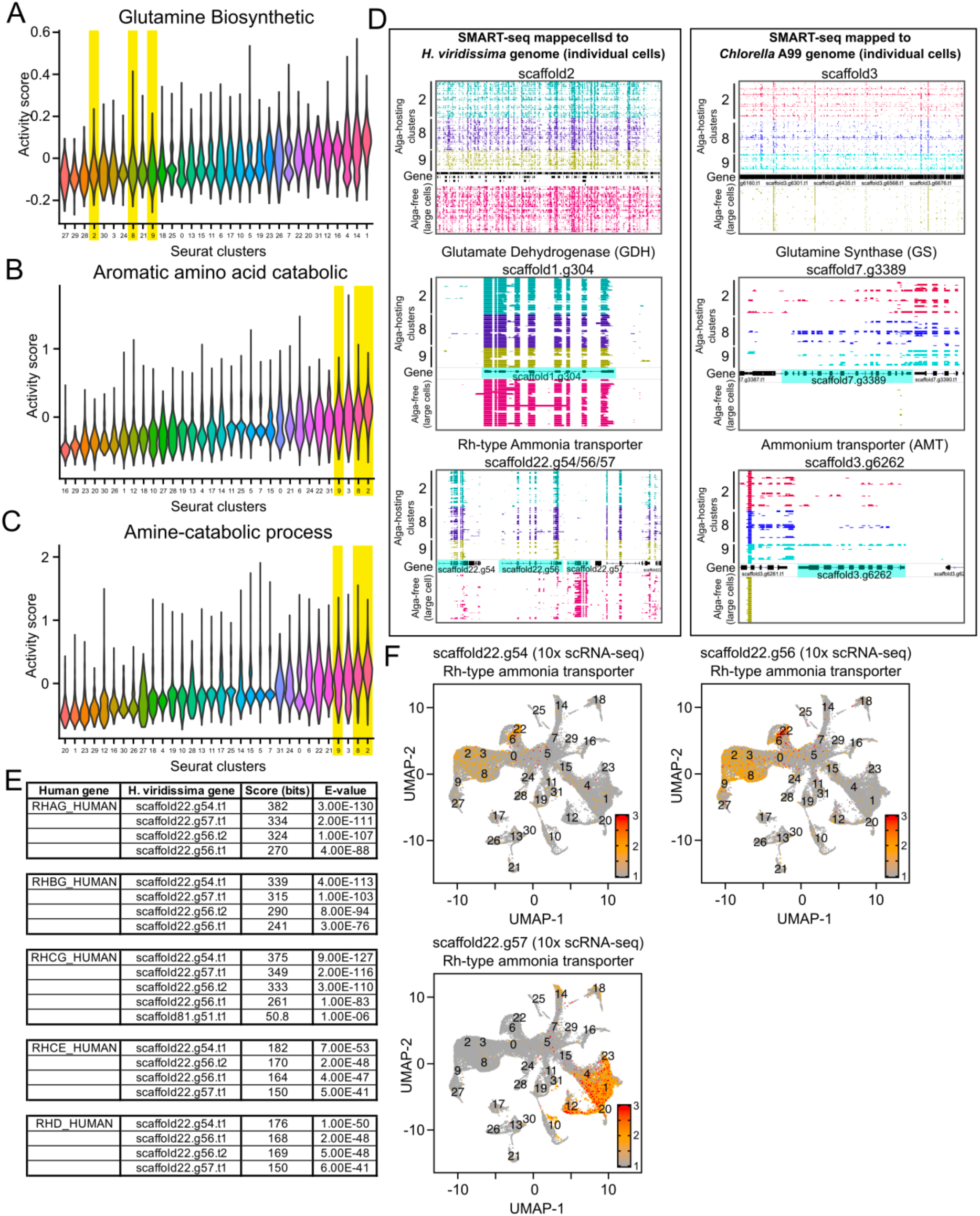
Nitrogen metabolism-related gene expression. **A-C.** Violin plot of activity scores for glutamine biosynthesis (**A**), aromatic amino acid catabolic (**B**) and amine catabolic process activity (**C**). The alga-hosting cell clusters (highlighted in yellow) are low in glutamine biosynthetic activity but high in activities related to aromatic amino acid and amine catabolism. **D.** IGV Genome browser tracks showing SMART-seq transcriptome expression from individual cells across the indicated genome regions. Alga-hosting cells (clusters 2, 8, 9) and large alga-free cells are indicated. The left panels: Top, SMART-seq data mapped to the *H. viridissima* genome and show the entirety of scaffold2. Middle, the gene encoding Glutamate dehydrogenase (GDH, scaffold1.g304). Bottom, a cluster containing genes encoding Rh-type ammonia transporters (scaffold22.g54, 56, 57). The right panels are SMART-seq data mapped to the *Chlorella* A99 genome. Top, the gene encoding for Glutamine Synthetase (GS, scaffold3). Middle, the gene encoding Glutamine Synthase (GS, scaffold7.g3389). Bottom, the gene encoding Ammonium transporter (AMT, scaffold3.g6262). The gene model is highlighted in cyan. **E.** Table showing homology (BLAST scores and E-values) between human Rh family glycoprotein ammonia transporters (RHAG, RHBG, RHCG, RHCE, RHD) and the identified putative Rh-type ammonia transporter genes scaffold22.g54, scaffold22.g56 and scaffold22.g57 from *H. viridissima*. **F.** UMAP plots illustrating the expression of *H. viridissima* Rh-type ammonia transporter gene homologs scaffold22.g54, scaffold22.g56, and scaffold22.g57 across all cell clusters in the 10x scRNA-seq dataset.

**Figure S4.**
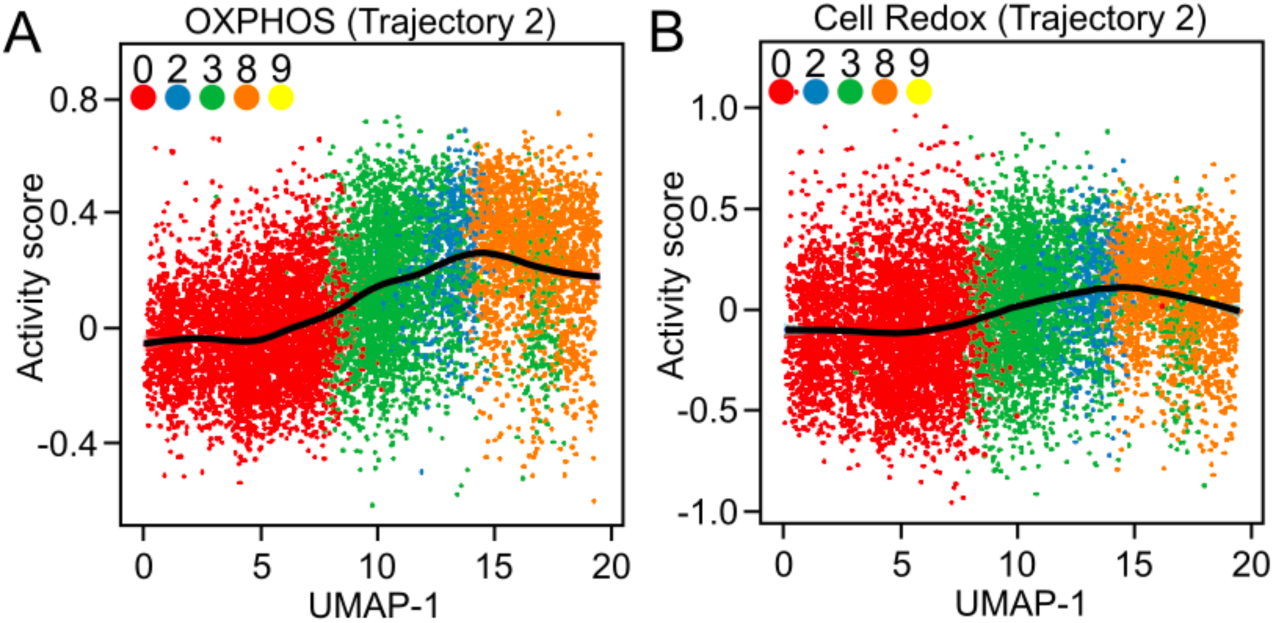
Lineage trajectory analyses and OXPHOS and redox activities along the trajectory. **A. B.** Plot showing the OXPHOS activity (**A**) and cell redox activity (**B**) along pseudo-time for lineage 2.

**Figure S5.**
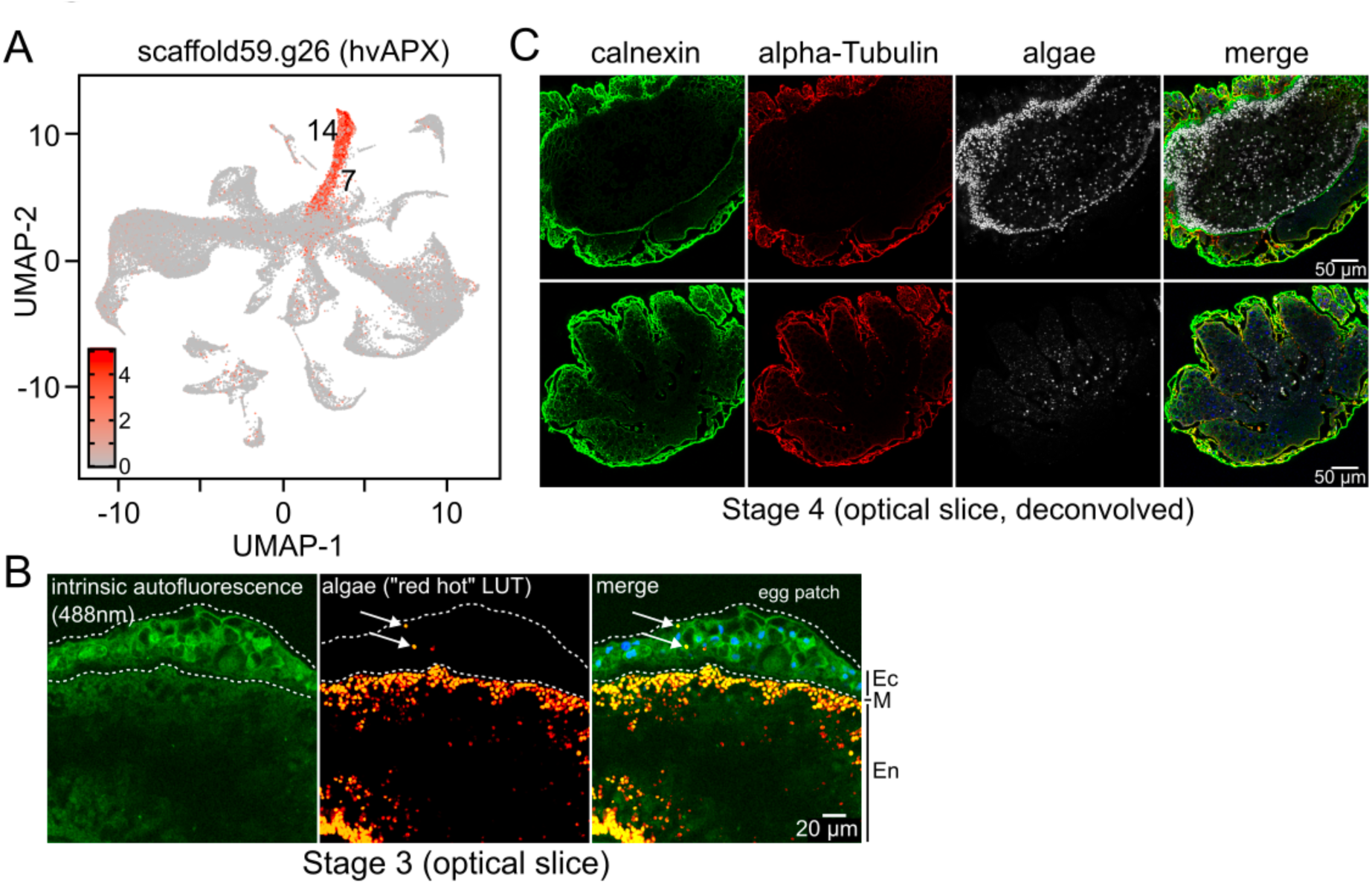
Gene expression in nurse cells and oogenesis samples. **A.** UMAP plot showing the expression of *H. viridissima* Ascorbate Peroxidase (hvAPX, scaffold59.g26), a plant-derived peroxidase, predominantly in nurse cell clusters. **B.** Confocal microscopy image of a cleared stage 3 egg-patch bearing Hydra (optical slice). The intrinsic autofluorescence of the whole mount is presented in green (“intrinsic autofluorescence 488nm”), algae in “red hot” LUT, and a merge. DAPI staining is in cyan but not presented as a separate panel. White arrows point to algae within the stage 3 egg-patch. The ectoderm (Ec), mesoglea (M) and endoderm (En) are denoted by black bars on the outside of the image. Scale bar = 20 µm. **C.** Confocal microscopy images of a cleared stage 4 oocyte (optical slices, deconvolved) stained for Calnexin (cell-boundaries, green), alpha-tubulin (red), and algal autofluorescence (greyscale). The top panel shows details deeper into the egg-patch region, and into the endoderm, while the bottom panel shows details within the oocyte, which is on the surface. Scale bar = 50 µm.

## Materials and methods

### Animal husbandry

*Hydra viridissima* were maintained in plastic tanks containing 1 L of artificial *Hydra* medium (0.5x ABC). The 0.5x ABC medium was prepared by adding 2 mL each of three (“A”, “B”, and “C”) concentrated stock solutions to 2 L of deionized water under continuous stirring. Stock “A”: 14.61 g NaCl, 1.864 g KCl and 27.745 g CaCl₂ in 500 mL deionized water; Stock “B”: 30.285 g Tris base in 500 mL deionized water. Stock “C”: 3 g MgSO₄ in 500 mL deionized water. The 1L tanks were housed in a Percival incubator set to 25 °C with a 12/12-hour light/dark cycle. *Hydra* were fed Artemia (San Francisco strain) daily. To enrich for animals with egg patches, the *Hydra* were grown at high density and only fed twice a week on Tuesday and Thursday. *Hydra* were transferred to clean tanks using a transfer pipette every one or two weeks, or when culture conditions deteriorated. Care was taken to avoid transferring debris such as dead Artemia or waste. A plastic pipette was used when necessary to gently dislodge and collect animals. To standardize physiological state, *Hydra* were transferred to fresh medium and starved for 48 h prior to all experiments.

### Tissue dissociation, single-cell preparation and 10x scRNA-seq

For scRNA sequencing, 80–100 *Hydra* viridissima polyps were transferred to a 1.5 mL microcentrifuge tube and washed twice with fresh 0.5x ABC *Hydra* medium. Following removal of the supernatant, animals were dissociated into a single-cell suspension by incubation in 15 mg/mL Pronase E (Roche), dissolved in 0.1× phosphate-buffered saline (PBS), for ∼20 min at room temperature with occasional gentle pipetting to facilitate tissue breakdown. Upon completion of dissociation, 0.01% bovine serum albumin (BSA, Sigma) was added. The cells were centrifuged at 170g for 5 min at 4 °C, and the resulting pellet was gently resuspended in 0.1× PBS and washed twice to ensure the removal of residual enzymes and debris. The final suspension was filtered twice through a 40 µm nylon mesh to eliminate tissue fragments. For the oogenesis sample, tissue surrounding the egg patch was microdissected and region containing the egg-patch was dissociated using the same protocol. Approximately 17,000 cells per sample were used for single-cell library preparation using the 10x Genomics Chromium Single Cell 3′ v3 or v4 platform, following the manufacturer’s instructions.

### Fluorescence-assisted cell sorting (FACS) of dissociated Hydra cells

Dissociated *Hydra* cells as described above in “Tissue dissociation and single-cell preparation” were filtered through a 40 µm nylon mesh prior to staining with Hoechst 33342. Staining was done for 20 min at room temperature. FACS was done on either BD Aria III or a BD Symphony using a 100 µm nozzle and cells were deposited into tubes containing 0.5x ABC. FACS profiling and gating setup was done using DAPI and PerCP-Cy5.

### SMART-seq and Bulk RNA-seq

Dissociated *Hydra* cells as described in “Tissue dissociation and single-cell preparation” were filtered through a 40 µm nylon mesh and chilled on ice. Cells were resuspended in 0.5x ABC containing 1% BSA. Serial dilutions of cells were then prepared on a 10 cm dish in 0.5x ABC containing 1% BSA. Individual cells were picked by mouth-pipette under a bright-field or TE200 invert fluorescence microscope with a 30 µm glass needle. Algae content was verified by imaging the isolated cell on the TE200 microscope using the Cy5 channel prior to transferring to SMART-seq lysis solution. To avoid premature cell lysis, we limited cell picking to a maximum of 30 minutes after cell preparation, and accumulated cells over multiple sessions. SMART-seq cDNA and library preparation (Takara, R400750) was done according to the manufacturer’s instructions.

### RNA preparation and sequencing

RNA for bulk RNA-seq was extracted using Trizol and the Zymo Direct-Zol kit. Library production for bulk RNA-seq was done using the Illumina TruSeq Stranded mRNA (Illumina, 20020594) platform according the manufacturer’s instructions. Nucleic acids were quantified using the Qubit 3 fluorometer and their profiles were examined using an Agilent Bioanalyzer. Sequencing was done using SBS-XLEAP P2 flow cells on the Illumina NextSeq 1000 platform.

### Mapping of single-cell transcriptomic libraries

Reads from scRNA-seq libraries were mapped to the *Hydra viridissima* genome (v1, from https://marinegenomics.oist.jp/hydra_viridissima_a99/viewer/download?project_id=82) using CellRanger to obtain counts of unique molecular identifiers (UMI) per gene and cell. Seurat objects were created in R with Read10X and CreateSeuratObjectfunction from Seurat (v.4.0.3). The filtered_feature_bc_matrix in the CellRanger output is used as data.dir for the Read10X function. Cells with fewer than 200 genes were excluded. The non-oogenesis samples were integrated into a single Seurat object with the IntegrateData function. Clusters were defined with FindClusters with a resolution of 0.6. Marker genes for each cluster were identified with FindAllMarkers. Activity scores for different gene modules are calculated by AddModuleScore function. To annotate gene functions, we identified the best SwissProt hit for each gene and assigned the corresponding GO terms from that hit to the input gene. This allowed us to extract genes associated with specific biological functions. For example, we selected all genes annotated with phagocytosis (GO:0006909), glutamine biosynthesis (GO:0009084) and ECM disassembly (GO:0022617) for module activity analysis. For other modules, we first performed GO enrichment analysis on Cluster 2 marker genes, then used all marker genes associated with the enriched GO terms to assess activity across all clusters. The gene lists used for activity score calculations are provided in Table S1. The similarity between each cluster and the cell types defined in the *Hydra vulgaris* were calculated by get_mapping_scores from SAMap (v. 0.1.4). The most similar biological cell type in *Hydra vulgaris* is used to defined the biological cell type for each of the *Hydra viridissima* cluster. For SMART-seq, the most similar cluster from Hydra viridissima was used Table S2.

### Trajectory inference analysis

Trajectory analysis was performed using the Slingshot (v1.8.0) to reconstruct pseudotemporal relationships among cell clusters. A subset of cells representing symbiotic lineages was extracted from the integrated dataset by selecting clusters 5, 0, 3, 2, 8, and 9, based on preliminary UMAP and clustering results. To designate a biologically meaningful origin, the interstitial stem cell cluster (iSC, cluster 5) was specified as the root of the trajectory. UMAP dimensionality reduction was performed using the top 20 principal components, and UMAP embeddings were manually assigned to the reducedDims slot of a SingleCellExperiment object. Slingshot was then applied to the UMAP embedding using cluster identities as labels and the starting cluster defined as cluster 5. An additional analysis excluding the interstitial stem cell population (clusters 0, 3, 2, 8, and 9) was also conducted with cluster 0 as the starting point. A pseudotime value is assigned to each cell and used for analysis of activity score along developmental trajectory.

### Fluorescence in situ hybridization (FISH)

Gene expression was visualized using RNA fluorescence in situ hybridization (FISH). A select gene transcript was detected using the single-molecule inexpensive FISH (smiFISH) protocol as previously described[47]. The gene-specific probe sets, available in Table S3, designed against the scaffold35.g35 transcript and labeled with fluorophore-conjugated secondary oligonucleotides. Due to the inability to detect scaffold1.g408 using smiFISH, a more sensitive detection method was employed based on a modified version of the PHYTOMap protocol[48]. *Hydra viridissima* were first relaxed in 2% urethane for 1 min, then fixed in 1% formaldehyde in *Hydra* 0.5x ABC (973 μL 0.5x ABC and 27 μL 37% formaldehyde) for 30 min at room temperature. Samples were transferred to ice-cold methanol (–20 °C) for 5 min, followed by three 10 min washes in RNase-free PBS. Samples were then dehydrated overnight in 30% sucrose at 4 °C, followed by incubating for 2-h at 4 °C in a 1:1 mixture of 30% sucrose and O.C.T. Compound, embedded in OCT, and snap-frozen in ethanol/dry ice. Tissue sections (10 μm) were cut and stored at –80 °C. Before hybridization, slides were air-dried at room temperature for 3–5 min and fixed in 1% formaldehyde for 5 min, followed by three PBS washes (3 min each). Sections were permeabilized with Proteinase K (20 mg/ml, 1:2000 dilution in PBSTR (PBS with 0.1% Tween-20 and 1:100 RNase inhibitor)) for 8 min at room temperature, followed by two 3 min washes with PBSTR. Hybridization was carried out using using three sets of scaffold1.g408-specific SNAIL padlock and SNAIL primer probes (500 nM per oligo) pre-annealed at 90 °C for 3 min and cooled to room temperature. The hybridization mixture contained 2× SSC, 30% formamide, 1% Triton X-100, 1% RNase inhibitor, and 10 nM probe mix in DEPC-treated water. Tissues were incubated at 40 °C for 3 h or overnight. Post-hybridization washes included two 20 min washes with PBSTR at 37 °C and one 30 min wash in 4× SSC with PBSTR at 37 °C, followed by a 1 min rinse in PBSTR at room temperature. Ligation was performed by first incubating sections on ice in ligation buffer (10× ligation buffer, 2 mg/mL BSA, RNase inhibitor, and DEPC-treated water) for 5 min without ligase, followed by a 4 h incubation at room temperature with 2 μL T4 DNA ligase added. For rolling circle amplification (RCA), tissues were preincubated on ice for 5 min in RCA mix without polymerase, followed by overnight incubation at 30 °C with equiPhi29 DNA polymerase (Lucigen). Following RCA, tissues were washed twice in PBSTR (10 min each), then twice in 2× SSC (1 min each). Detection was performed using fluorescent detection oligonucleotides (100 nM per probe) in hybridization buffer (20× SSC, 20% formamide, and DEPC-treated water) for 1 h at room temperature. Final washes include two 5 min rinses in 2× SSC and DAPI staining (1 mM, 2 min). Samples were washed twice in DEPC-treated water and mounted with antifade mounting medium. Confocal imaging was performed using a Zeiss LSM microscope

### Fixation of dissociated cells

Dissociated cells, as described in the section “Tissue dissociation and single-cell preparation” were fixed by adding 16% paraformaldehyde (PFA; EMS, 15710) to a final of 4% and incubating for 10 min at room temperature. The fixative was neutralized with 125 mM glycine for 5 minutes at room temperature.

### Fixation of *Hydra viridissima* for whole mount staining

*Hydra* were collected in a single well of a 24-well plate containing 0.5x ABC. The 0.5x ABC was removed and replaced with 0.5x ABC containing 2% urethane for 1-2 min. 16% PFA (EMS, 15710) was added to a final of 4% and incubated for 10 min at room temperature. After 10 min, the fixative was aspirated and replaced with 4% PFA in 0.5x ABC. The *Hydra* were incubated in fresh fixative for another 20 min. The fixative was neutralized with 125 mM glycine for 5 min at room temperature.

### Surface biotinylation of dissociated cells

PFA-fixed cells from “Fixation of dissociated cells” were pelleted at 500 g x 5 min at room temperature and washed twice with 0.1x PBS. The final cell pellet was resuspended in 100 μl of 1x PBS, 8 μl of a 10 mM stock solution of Sulfo-NHS-SS-Biotin (Thermo, A39258) was added (final concentration: 0.74 mM) and incubated at room temperature for 30 min. The reaction was quenched with 1M Tris-HCl pH 8 at a final concentration of 25 mM. The cells in neutralized reaction solution were then pelleted at 500 g for 5 min and washed with 1x PBS. The cells were resuspended in 1x PBS and used for Streptavidin staining.

### Peroxidase activity in whole mount *Hydra*

Fixed *Hydra* from “Fixation of Hydra for whole mount” step were first permeabilized with PBS containing 0.25% Triton X-100 (PBSTx) for 10 min at room temperature. The solution was aspirated and the *Hydra* were washed with 1x PBS. A 1 ml solution of Biotin-phenol (Iris Biotech; LS-3500.0250) at 125 μM was next prepared in 0.5x ABC. 500 μl of this solution was mixed with 200 μl of McCoy’s 5a media containing 10% FBS to make the reaction solution. The reaction solution was then added to *Hydra*, after aspirating the PBS, and incubated at room temperature with gently rocking for 30 min. 2 μl of a 100 mM Hydrogen Peroxide solution was added to the solution, mixed by gently flicking and incubated at room temperature for 10 min. The reaction was halted by first aspirating off the solution and then washing with 1x PBS containing 10 mM sodium ascorbate and 10 mM Trolox. The *Hydra* were then washed with 1x PBS.

### Streptavidin staining of surface biotinylated dissociated cells

Fixed cells from “Peroxidase activity in whole mount Hydra” were first permeabilized with 1x PBS containing 0.25% Triton X-100 (PBSTx) for 30 minutes at room temperature. Cells were centrifuged at 500 g x 5 min and the permeabilizing solution was aspirated. The cells were then blocked in Streptavidin Blocking solution (SBS), which is 1x PBS containing 0.1% Tropix i-Block biotin-free casein, 0.1% Tween-20 and DAPI (1 μg/ml) for 30 min at room temperature. Alexa-fluor conjugated Streptavidin was prepared in SBS at 1:300. Cells were pelleted at 500 g x 5 min and aspirated. The cells were resuspended in SBS containing Streptavidin, and incubated for 1 h at room temperature. The cells were pelleted at 500 g x 5 min and washed 3 for 5 min each with 1x PBS containing 0.1% Tween-20. The final cell pellet was resuspended in 5 μl of 1x PBS plus 0.1% Tween-20. 10 μl of Prolong antifade gold was added and mixed and spotted on SuperFrost Plus microscope slides for viewing.

### Streptavidin staining of whole mount Peroxidase activity

*Hydra* from the “Peroxidase activity in whole mount Hydra” step were blocked in Streptavidin Blocking Solution (SBS, 1x PBS containing 0.1% Tropix i-Block biotin-free casein (Applied Biosystems, T2015), 0.1% Tween-20 and DAPI (1 μg/ml)) for 30 min at room temperature. Alexa-fluor conjugated Streptavidin was prepared in SBS at 1:300. The *Hydra* were resuspended in SBS containing Streptavidin, and incubated for 1 h at room temperature. The staining solution was aspirated and the *Hydra* washed 3 x 5 min each with 1x PBS containing 0.1% Tween-20. Mounting and tissue clearing was done as described below in the “Whole mounting and tissue clearing” step. In cases where immunostaining was combined with Streptavidin, the “Immunostaining of whole mount Hydra” procedure was done first, and the Streptavidin was included in the secondary antibody step.

### Immunostaining of whole mount *Hydra*

Fixed *Hydra*, as described in “Fixation of Hydra for whole mount”, were first permeabilized with 1x PBS containing 1% SDS for 15 min. The SDS solution was aspirated and 1x PBS containing 0.25% Triton X-100 for 30 min at room temperature. The *Hydra* were then blocked in Antibody Blocking Solution (ABS), which is 1x PBS containing 10% normal goat serum, 10% BSA, 0.1% Tween-20 and 10 mM sodium azide for 30 min at room temperature. Antibodies were prepared in ABS at the dilutions indicated in Table S4. *Hydra* were pelleted at 500 g x 5 min and aspirated. The *Hydra* were resuspended in ABS containing antibodies, and incubated overnight (up to two days) at room temperature in a humidified chamber. The next day, the *Hydra* were pelleted at 500 g for 5 min and washed 3 x 5 min each with 1x PBS containing 0.1% Tween-20. *Hydra* were aspirated and resuspended in secondary antibodies that were prepared in 1x PBS containing 0.1% Tween-20 for 1-2 h. DAPI at a final of 1 μg/ml was added to the secondary antibody solution. *Hydra* were pelleted at 500 g for 5 min and washed 3 x 5 min each with 1x PBS containing 0.1% Tween-20. The stained *Hydra* were resuspended in 1x PBS plus 0.1% Tween-20. Mounting and tissue clearing was done as described below in the “Whole mounting and tissue clearing” step.

### Whole mount and tissue clearing

To image whole mount Hydra, two 1.5 coverslips were attached to a Superfrost Plus slide with nail polish to form a channel. The stained *Hydra* from the above were then deposited into the channel formed by the 1.5 coverslips, positioned, and then the solution was aspirated. The Ce3D tissue clearing reagent (Biolegend, 427701), supplemented with DAPI (1 μg/ml) was added to submerge the *Hydra*, and incubated until the tissue was clear. The CE3D tissue clearing reagent was then removed and Prolong antifade gold supplemented with DAPI (1 μg/ml) was added. The stained *Hydra* were then covered with a #1 coverslip and sealed with nail polish for imaging. We note that the tissue clearing agent used here results in a pronounced fading of the DAPI signal within one or two days. We imaged the whole mount sample the within one or two days.

### Microscopy

Wide-field images were acquired using MetaMorph software with a 10x (NA 0.3) objective on a Nikon E800 equipped with an ORCA Flash 4.0 LT camera. Confocal imaging was done using Leica AF software with a 40x (NA 1.3) oil objective on a Leica SP5 scanning confocal microscope. Images were processed with FIJI/ImageJ version 2.14.0/1.54f using “Enhance contrast” and “bleach correction”. Deconvolution was done with the DeconvolutionLab2 2.1.2 (27.06.2018) plugin[49].

## Data Availability

All sequencing data is available at NCBI GEO accession number GSE306770

**Table S1. Genes used for predicting gene module activity in the scRNA-seq dataset. Each sheet is named after the corresponding gene module.**

**Table S2. SMART-seq cell classifications and cluster prediction.**

**Table S3. Oligos used for Fluorescence in situ hybridization.**

**Table S4. Antibodies and staining reagents used in this study.**

**Video S1. Algae found in the stage 3 egg patch.**

**Video S2. Alpha-tubulin process transverse the stage 4 oocyte from the ectoderm to the mesoglea.**

**Video S3. Alpha-tubulin process run along the mesoglea.**

**Video S4. Alpha-tubulin process merge to form a “bouquet”-like structure and are associated with algae**

**Video S5. Imaging of tissue cleared stage 5 egg-patch reveals mesoglea structure (calnexin), tubulin processes (alpha-tubulin) and the presence of algae within the oocyte.**

**Video S6. The stage 5 oocyte shows cell processes contacting the mesoglea that can contain algae.**

